# Factors associated with the use of maternal health services by mothers in a post-conflict area of western Côte d’Ivoire in 2016

**DOI:** 10.1101/673970

**Authors:** Samba Mamadou, Attia-Konan Akissi Régine, Sangaré Abou Dramane, Youan Gotré Jules, Kouadio Luc Philippe, Bakayoko-Ly Ramata

## Abstract

**Introduction:** In Côte d’Ivoire, maternal health service utilization indicators remain low despite improvements in health coverage and the availability of free health care for pregnant women. The objective of the study was to identify the determinants associated with the use of maternal health services in the department of Bloléquin, in western Côte d’Ivoire.

**Methods:** We conducted a cross-sectional study with an analytical focus. The study sample size was 400 women. Study participants were selected through a two-stage cluster survey. The data were collected using a standardized questionnaire whose items concerned socio-demographic data, the different uses of maternal health services, namely childbirth assisted by a health professional, use of family planning, prenatal consultation and post-natal consultation. Logistic regression was used to investigate factors associated with the use of maternal health services. The significance of the statistical tests was set at 5%. The odds ratios and 95% confidence intervals were calculated and interpreted.

**Results:** The results showed that women made less use of family planning services (OR=0.4), prenatal consultation (OR=0.2) and assisted childbirth (OR=0.2) when they provided the funding for care themselves. Women with incomes above $26.8 used family planning services 4 times more than those with lower incomes. Married women used prenatal consultations 3 times more often than unmarried women (CI_95%_ = 1.4 - 7.3). Desiring pregnancy increased the use of post-natal consultations by 3 times (CI_95%_ = 1.5 - 6.1).

**Conclusion:** Improving the use of maternal health services in western Côte d’Ivoire requires taking into account women’s socio-cultural and economic challenges. In initiatives related to the financial empowerment of women, efforts must be made at the level of emotional factors related to pregnancy.

## Introduction

Access to maternal health services is a challenge for health systems in developing countries. In fact, the use of these services makes it possible to reduce obstetric complications, thus maternal mortality. Although the number of maternal deaths decreased from 532,000 to 303,000 between 1990 and 2015 worldwide, maternal mortality remains the highest in sub-Saharan Africa with a rate of 546 deaths per 100,000 live births [1]. To reduce this rate, most states in Africa have introduced an exemption from the cost of care for pregnant women [2]. However, maternal health services are underutilized, especially in low- and middle-income countries.

In Côte d’Ivoire, indicators of maternal health service utilization are low despite the existence of a national policy of “free care” for pregnant women. The coverage rate in CPN4 was 38.23% in 2015, the modern contraceptive prevalence rate was 17%, and the assisted childbirth rate was 55.10% [3]. The maternal mortality ratio was estimated at 645 per 100,000 live births, well above the regional average [1]. The western part of the country, which was particularly affected by the military-political crisis between 2002 and 2011, is experiencing alarming rates. There is significant underutilization of maternal health services with a 25% ANC 4 rate compared to 35% for the national average, with high ANC drop-out rates of up to 67% [3]. To improve access to care, the Government has introduced a policy of free care for pregnant women, taking into account prenatal consultations (ANC), childbirth and caesarean section throughout the country. In addition, there has been an increase in the coverage of health facilities. However, maternal health services remain underutilized and the factors associated with this underutilization by women are not well known, although the literature describes some of them, including economic, social, cultural and organizational parameters related to the health system [4]. A good knowledge of these determinants will certainly make it possible to better address the issue of the use of maternal health services. This is why it seemed wise to understand the social, cultural and economic logic of the use of maternal care of people living in this part of Côte d’Ivoire The objective of the study was to identify the determinants associated with the use of maternal health services in the department of Blolequin, in western Côte d’Ivoire

## Methods

### Study area

The transversal analytical study took place in the department of Blolequin located in the West of Côte d’Ivoire. It has one general hospital, two urban health centres and five rural health centres at the public health level. In 2016, the number of health workers was estimated at 36, including 7 general practitioners, 19 nurses and 10 midwives, for an estimated population of 103,189 inhabitants and 21,324 women of childbearing age.

### Sampling and setting

The study population consisted of women with a child aged up to 3 years old at the time and living in the Blolequin department at the time of the survey. Were included in the study sample, women meeting these inclusion criteria and giving their informed consent. The sample size was calculated from Schwartz’s formula:

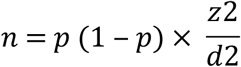

with p = 38% (corresponding to the national level of ANC 4 rate according to the 2013 RASS); z = 1.96; d = 0.05; n=362. By increasing this number by 10%, the sample size was 400 women.

For the recruitment of women to be interviewed, two-stage cluster sampling was used. The eight sanitary areas of the department formed the clusters. In the first degree, six areas were drawn at random. Then, in the second degree, a random draw of two secondary roads in each sanitary area was carried out.

### Data collection

The data was collected by 4 investigators trained under the supervision of a lead investigator Once in the chosen locality, the direction to be taken by the investigator was determined by the bottle method. Thus, once in the street drawn at random, the direction was indicated by the direction taken by the tip of the bottle following a throw. Then a random draw for a concession from a random number table was used to find the first household to survey. The survey was carried out from one person to the next until the number of women required per area was reached. The number of women required per area was allocated in proportion to the number of women of childbearing age in each area. The data were collected using a standardized questionnaire whose items related socio-demographic, economic, maternal health practices and constraints related to geographical and financial accessibility to health facilities.

### Data analysis

A descriptive analysis of the variables was carried out using frequencies and percentages. A univariate analysis tested the association between the variables of interest and each of the explanatory variables. This step identified the variables to be included in the multivariate analysis. The significance threshold was set at p < 20%. Logistic regression was used to investigate factors associated with the use of maternal health services.

There were four variables of interest related to service utilization: childbirth attended by a health professional, family planning (FP), prenatal consultation (PN), and post-natal consultation (PNC). The independent variables included age, marital status, household type, place of residence, gestational age, desire for pregnancy, estimated income, source of funding, distance to a health centre. The final model of the logistic regression was obtained by the top-down step-by-step method. The odds ratios were calculated to measure the strength of association between the variables studied. The significance of the statistical tests was set at 5%. The data collected were analysed by the STATA 12.1 software. The authorizations of the administrative and health authorities of the department of Blolequin were obtained as well as the verbal informed consent of the women surveyed. The confidentiality of the data was preserved throughout the study.

### Ethical considerations

The study protocol was approved by the National Research Ethics Committee of the Ministry of Health and Public Hygiene of Côte d’Ivoire.

## Results

### Sociodemographic and economic characteristics

Women of reproductive age were predominantly young and lived in couples (85.5%). Nearly half of the women were between the ages of 22 and 30 years old. They were mostly housewives (47%) and had no regular income (36.5%). They lived in cities less than 5 km from a health facility (75%). Women lived 75% in a monogamous home. The spouse provided care financing for 80% of the women (Table 1).

**Table 1:** Socio-demographic and economic characteristics of the sample (N = 400)

| Variables |  | Frequency | Percentage (%) |
| --- | --- | --- | --- |
| Age (year) | [16-22[ | 99 | 24,7 |
|  | [22-31[ | 178 | 44,5 |
|  | [31-49] | 123 | 30,8 |
| Marital status | Unmarried | 32 | 8,0 |
|  | Married | 342 | 85,5 |
|  | Widowed/Divorced | 26 | 6,5 |
| Professional activities | Housewife | 188 | 47,0 |
|  | Farmer | 76 | 19,0 |
|  | tradeswoman | 52 | 13,0 |
|  | Craftsman | 48 | 12,0 |
|  | State agent | 24 | 6,0 |
|  | Other | 16 | 4,0 |
| Monthly income (US\$) | No income | 146 | 36,5 |
|  | <26,8 | 78 | 19,5 |
|  | 26,8 - 53,5 | 85 | 21,3 |
|  | >53,5 | 91 | 22,8 |
| Type of relationship | Polygamous | 101 | 25,3 |
|  | Monogamous | 299 | 74,7 |
| Place of residence | Urban | 234 | 58,5 |
|  | Rural | 166 | 41,5 |
| Distance to health facility | ≤5 km | 300 | 75,0 |
|  | >5 km | 100 | 25,0 |
| Funding source | Spouse | 322 | 80,5 |
|  | Personal income | 45 | 11,3 |
|  | Parents | 33 | 8,2 |
| Desired pregnancy | Yes | 354 | 88,5 |
|  | No | 46 | 11,5 |
| Gestated | 1 | 80 | 20,0 |
|  | >1 | 320 | 80,0 |

### Frequency of use of maternal health services

Family planning services are the least (40%), followed by ANCs (58.8%). Assisted childbirth and PNCs were the most commonly used, with respectively 78.5% and 78.25%, or about 8 out of 10 women (Table 2).

**Table 2:** Frequency of use of maternal health services (N=400)

| Type of service | Frequency | Percentage (%) |
| --- | --- | --- |
| Family planning (FP) | 160 | 40,0 |
| Prenatal consultations (ANC) | 235 | 58,8 |
| Assisted childbirth (PNC) | 314 | 78,5 |
| CPON | 313 | 78,3 |

### Determinants of maternal health service utilization

#### Family planning services

Women with monthly incomes of at least $ 26.8 use more family planning services than women with no income (OR = 4.0 and OR = 3.9). When women self-financed care, they used family planning services less (OR = 0.4) (Table 3).

**Table 3:**
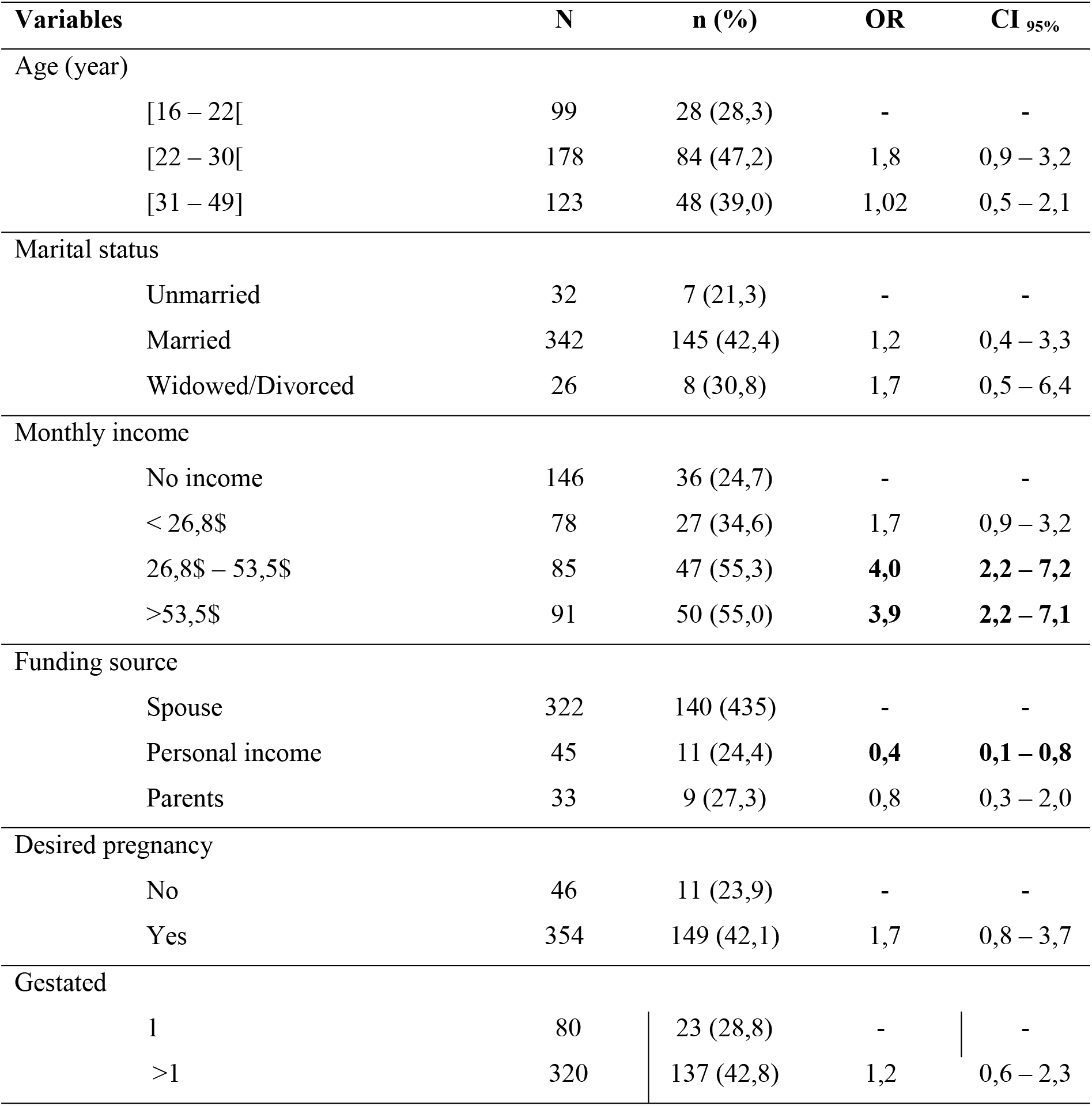
Determinants of family planning service utilization in Blolequin department: logistic regression (N=400)

### Prenatal consultations service (ANC Services)

Marital status and source of funding significantly influence the use of ANC services. Married women used ANC services more than unmarried women (OR=3.2). Those who financed their own health expenses used ANC services less (OR = 0.2) (Table 4).

**Table 4:** determinants of the use of ANC services (N = 400)

| **Variables** | **N** | **n (%)** | **OR** | **IC<sub>95%</sub>** |
| --- | --- | --- | --- | --- |
| Marital status |  |  |  |  |
| Unmarried | 32 | 10 (31,3) | - | - |
| Married | 342 | 214 (62,6) | **3,2** | **1,4 – 7,3** |
| Widowed/Divorced | 26 | 11 (42,3) | 1,5 | 0,5 – 4,7 |
| Distance to health facility |  |  |  |  |
| ≤5 km | 300 | 168 (56) | - | - |
| >5 km | 100 | 67 (67) | 1,6 | 0,9 – 2,6 |
| Funding source |  |  |  |  |
| Spouse | 322 | 189 (80,4) | - | - |
| Personal income | 45 | 27 (60) | **0,2** | **0,1 – 0,5** |
| Parents | 33 | 19 (57,6) | 0,5 | 0,2 – 1,2 |
| Desired pregnancy |  |  |  |  |
| No | 46 | 17 (37,0) | - | - |
| Yes | 354 | 218 (61,6) | 1,9 | 0,9-3,8 |

### Delivery assisted by qualified personnel

Women in couples, as well as widows and divorced women, knew that assisted childbirth was almost 4 times more common than unmarried women (OR=3.6; OR=3.9). Women who self-supported their health expenses used less assisted childbirth (OR=0.2) (Table 5).

**Table 5:** Determinants of the use of assisted childbirth (N=400)

| Variables | N | n (%) | OR | CI<sub>95%</sub> |
| --- | --- | --- | --- | --- |
| Age (year) |  |  |  |  |
| [16 - 22[ | 99 | 71 (71,7) | - | - |
| [22 - 30[ | 178 | 149 (83,7) | 1,4 | 0,7-2,8 |
| [31 - 49] | 123 | 94 (76,4) | 0,9 | 0,4-1,7 |
| Marital status |  |  |  |  |
| Spouse | 32 | 13 (40,6) | - | - |
| Married | 342 | 284 (83,0) | **3,6** | **1,5-8,7** |
| / Widowed / Divorced | 26 | 17 (65,4) | **3,9** | **1,1-13,3** |
| Monthly income |  |  |  |  |
| No income | 146 | 111 (76,0) | - | - |
| < 26,8 | 78 | 57 (73,1) | 0,7 | 0,3-1,4 |
| 26,8 – 53,5 | 85 | 68 (80,0) | 0,9 | 0,4-2,0 |
| >53,5 | 91 | 78 (85,7) | 1,2 | 0,4-2,9 |
| Residence area |  |  |  |  |
| Urban | 166 | 137 (82,5) |  |  |
| Rural | 234 | 177 (65,4) | 2,0 | 1,0-4,0 |
| Funding source |  |  |  |  |
| Spouse | 322 | 271(84,2) | - | - |
| Personal income | 45 | 24(53,3) | **0,2** | **0,1–0,5** |
| Parents | 33 | 19(57,6) | 0,5 | 0,2–1,2 |
| Desired pregnancy |  |  |  |  |
| Non | 46 | 28 (80,7) | - | - |
| Oui | 354 | 286 (60,9) | 1,5 | 0,7-3,3 |
| Gestated |  |  |  |  |
| 1 | 80 | 57 (71,3) | - | - |
| > 1 | 320 | 257 (80,3) | 1,12 | 0,5-2,4 |

### Postnatal consultations services (PNC services)

Desiring pregnancy increased the use of CPONs by 3 times (CI _95%_ = 1.5 - 6.1). The fact of financing her own care leads women to consult less after childbirth (OR = 0.3; IC _95%_ = 0.1 - 0.5) (Table 6).

**Table 6:**
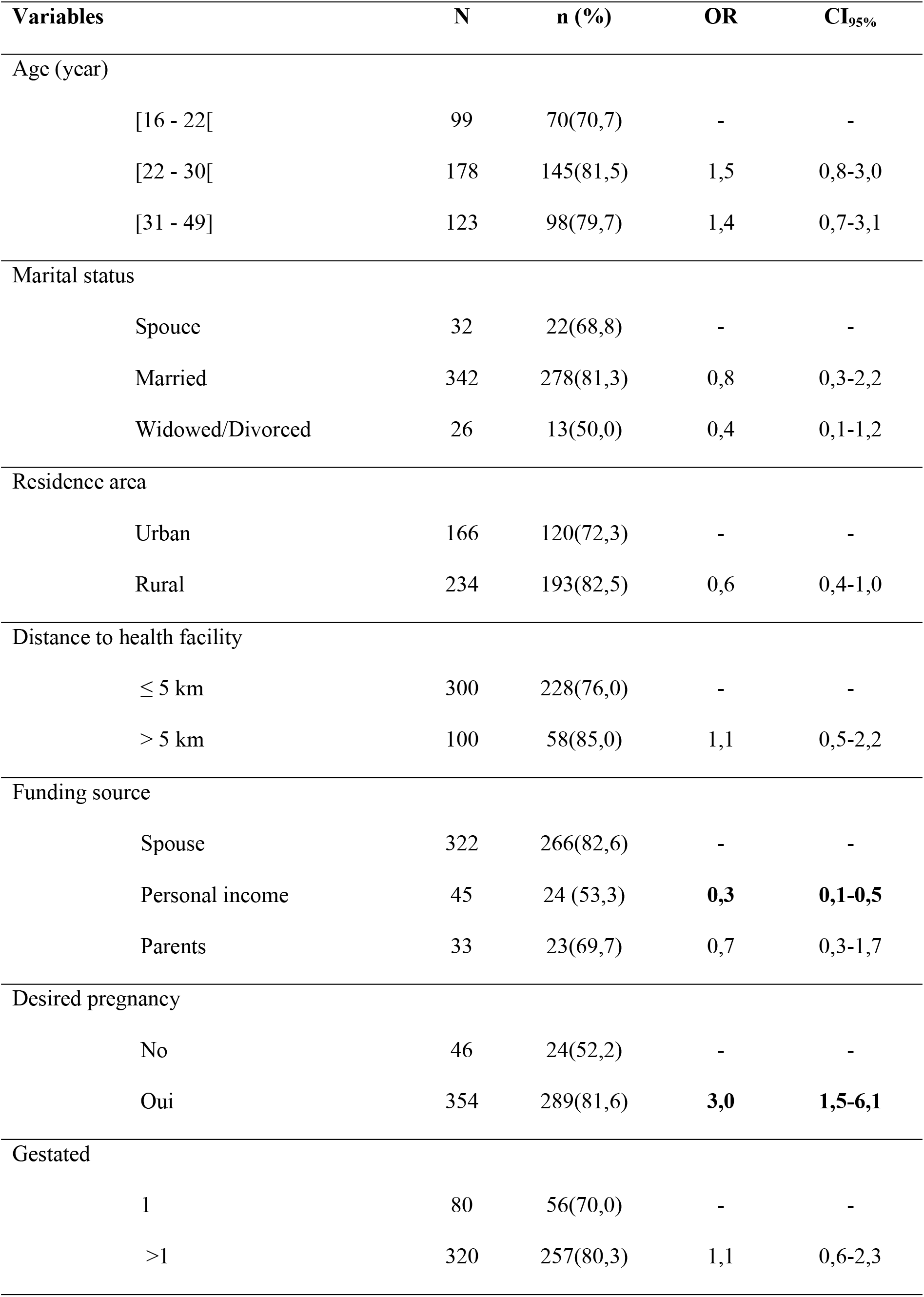
Determinants of the use of a post-natal consultation service (N=400)

| Variables | N | n (%) | OR | CI<sub>95%</sub> |
| --- | --- | --- | --- | --- |
| Age (year) |  |  |  |  |
| [16 - 22[ | 99 | 70(70,7) | - | - |
| [22 - 30[ | 178 | 145(81,5) | 1,5 | 0,8-3,0 |
| [31 - 49] | 123 | 98(79,7) | 1,4 | 0,7-3,1 |
| Marital status |  |  |  |  |
| Spouse | 32 | 22(68,8) | - | - |
| Married | 342 | 278(81,3) | 0,8 | 0,3-2,2 |
| Widowed/Divorced | 26 | 13(50,0) | 0,4 | 0,1-1,2 |
| Residence area |  |  |  |  |
| Urban | 166 | 120(72,3) | - | - |
| Rural | 234 | 193(82,5) | 0,6 | 0,4-1,0 |
| Distance to health facility |  |  |  |  |
| ≤ 5 km | 300 | 228(76,0) | - | - |
| > 5 km | 100 | 58(85,0) | 1,1 | 0,5-2,2 |
| Funding source |  |  |  |  |
| Spouse | 322 | 266(82,6) | - | - |
| Personal income | 45 | 24 (53,3) | **0,3** | **0,1-0,5** |
| Parents | 33 | 23(69,7) | 0,7 | 0,3-1,7 |
| Desired pregnancy |  |  |  |  |
| No | 46 | 24(52,2) | - | - |
| Oui | 354 | 289(81,6) | **3,0** | **1,5-6,1** |
| Gestated |  |  |  |  |
| 1 | 80 | 56(70,0) | - | - |
| >1 | 320 | 257(80,3) | 1,1 | 0,6-2,3 |

## Discussion

The use of maternal health services in the Blolequin department is low and far from the national average [3]. Assisted childbirth and postnatal consultations were the most commonly used services with similar rates. Po’s study showed a positive association between the three stages of obstetric care, ANC, childbirth and postnatal consultations (PNC) [5]. Hence the importance of setting up a monitoring reference system to ensure the continuity of services and thus increase the use of maternal health services.

Our study confirms the existence of a financial barrier to the use of maternal health services. Although maternal care is free in Côte d’Ivoire, there is still underutilization of maternal health services. A study conducted in Kenya, showed an increase in the use of maternal health services in cases of exemption measures in regions where there were public sector obstetric services available [6]. Hence the importance of increasing geographical accessibility to health centres. However, the financial barrier remains and women sometimes give up maternal care. When pregnant women funded care themselves, they used maternal health services less often. This is why the majority of them have proposed free care to improve its use. Indeed, most of the women in our sample were housewives and did not have a regular income. Several women indexed the quality of care, including reception and transportation costs, to a health facility. The provision of lower quality care has a negative impact on the use of care [7]. Transport-related expenses, which are not often taken into account in health expenditure, sometimes constitute an obstacle to the use of health services [8] even in the case of exemption from fees as observed in the in charge of tuberculosis in Burkina Faso [9]. Therefore, in addition to the obstacle of studies, they mentioned other obstacles to the use of maternal health services. These include mothers’ level of education, perception of care, lack of knowledge of obstetric complications and living in rural areas [10–11]. In addition, situations of political instability and armed conflict in a country compound the lack of access to maternal health services due to the deterioration of the health care system [12]. According to Gaber, the various socio-political crises that Côte d’Ivoire has experienced have had a negative impact on the supply of care [13].

Family Planning (FP) services were the least used among maternal health services despite the steps taken by the government to make FP products accessible with the help of development partners. FP services were more frequented by women with incomes. This suggests that having financial autonomy leads women to make a decision in favour of their health by avoiding multiple pregnancies. This result seems to contradict the fact that women used their income less to make prenatal consultations (ANC). He suggests that women pay more attention to family planning so they are willing to contribute financially. Nevertheless, it seems advisable to develop income-generating activities for these women in order to increase their financial autonomy. This action requires the popularization of literacy, although little cited by women as a means of improving the use of care (3 out of 400 women), as access to maternal care increases with the level of education [14]. However, women recognize that good awareness will promote better access, which can be increased through literacy. In fact, the use of maternal health services is increased when women have a better understanding of the number of recommended NPCs [15].

Marital status also influenced the use of maternal health services. Married women made greater use of ANC services and more use of assisted childbirth. Married women often benefit from the financial support of their spouses [15]. Single women who are pregnant use care less because they have fewer financial resources and do not always want to become pregnant [15]. They often leave the community to escape the accusatory gaze of society [16]. This phenomenon is magnified in girls although our study did not reveal age as an obstacle to the use of obstetric care. In fact, the latter often have a low rate of access to antenatal care [15; 17].

The level of acceptability of pregnancy impacts the use of PNC services. When women did not want to become pregnant, they used these services less. Beninguisse’s 2005 study identified this relationship but with ANC [19]. Thus, the opportunity of pregnancy appears to be a determinant of maternal care use. Therefore, the social dimension of the disease, especially when it is chronic or stigmatized, involves taking into account the psycho-emotional state of the patient to promote this use of care. This is what is observed in the management of HIV / AIDS. Although pregnancy is not a disease, it creates emotions for women that can influence their use of care. Improving well-being during pregnancy is therefore a concern for achieving the Sustainable Development Goals [20].

The parity and geographical access often associated with the use of obstetric care were not highlighted in our study [15]. The majority of women lived within 5 km of a health facility.

Our study presents the limitations of all observational studies regarding possible desirability information bias. Women may overestimate the frequency of use of maternal health services. Thus, we have higher reported rates than those calculated by the Ministry of Health, which collects data from health facility registers. This uniformly distributed information bias throughout the questionnaire was minimized by the administration of the questions by health workers with a good knowledge of the field.

## Conclusion

The underutilization of maternal health services is one of the causes of the high mortality rates observed in developing countries. The results of this study showed a positive association between the use of maternal health services, the marital status of women and the source of funding for care. Improving the use of maternal health services in the department of Blolequin involves taking into account these socio-cultural and economic challenges of women. Investing in the economic empowerment of women is a challenge in this area of Côte d’Ivoire that has been the scene of armed conflict. This autonomy will reduce the woman’s dependence and lead her to make informed choices about health. In addition to economic challenges, emotional factors of pregnancy need to be taken into account to improve this access. . The social dimension of pregnancy should be developed in maternal health services, where staff often only consider health aspects. Taking into account these different factors will contribute to achieving the objectives for sustainable development in Côte d’Ivoire.

## Author Contributions

**Conceptualization:** Samba Mamadou, Kouadio Luc Philippe, Bakayoko-Ly Ramata

**Methodology:** Samba Mamadou, Sangaré Abou Dramane, Attia Régine, Gotré Jules

**Investigation:** Gotre Jules

**Writing ± original draft:** Samba Mamadou, Sangare Abou Dramane, Attia Régine, Gotré Jules

**Writing ± review & editing:** Samba Mamadou, Kouadio Luc Philippe, Bakayoko-Ly Ramata, Sangaré Abou Dramane, Attia Régine, Gotré Jules

## Authors’ statement

The informed consent of the respondents was obtained prior to their participation in the study. Each study participant was briefed on the study objective and confidentiality was assured for any information provided. The authors do not declare any conflict of interest

